# Cross-platform normalization enables machine learning model training on microarray and RNA-Seq data simultaneously

**DOI:** 10.1101/118349

**Authors:** Jaclyn N. Taroni, Casey S. Greene

## Abstract

**Motivation:** Large compendia of gene expression data have proven valuable for the discovery of novel biological relationships. The majority of available RNA assays are run on microarray, while RNA-seq is becoming the platform of choice for new experiments. The data structure and distributions between the platforms differ, making it challenging to combine them. We performed supervised and unsupervised machine learning evaluations, as well as differential expression analyses, to assess which normalization methods are best suited for combining microarray and RNA-seq data.

**Results:** We find that quantile and Training Distribution Matching normalization allow for supervised and unsupervised model training on microarray and RNA-seq data simultaneously. Nonparanormal normalization and z-scores are also appropriate for some applications, including differential expression analysis.

**Availability and Implementation:** These analyses were performed in R and are available at https://www.github.com/greenelab/RNAseq_titration_results under a BSD-3 clause license.

*Supplementary Information* is available.

## 1. Introduction

The union of large and diverse compendia of gene expression data with machine learning approaches has enabled the extraction of cell type-specific networks (Greene et al., 2015) and the discovery of new biological patterns associated with cellular responses to the environment (Tan et al., 2016). Integrative analyses of multiple microarray cohorts have uncovered important signatures in human infection (Andres-Terre et al., 2015;Sweeney et al., 2016). Sequencing-based RNA assays have certain advantages over array-based methods, namely quantitative expression levels and a higher dynamic range (Wang et al., 2009). As a result, researchers have increasingly adopted this new technology for their gene expression experiments.

RNA-sequencing (RNA-seq) assays are a growing share of new gene expression experiments. In 2015, the ArrayExpress team reported that the amount of sequencing-based submissions had doubled in the last 18 months (Kolesnikov et al., 2015). However, it remains essential to include microarray data in large-scale studies of gene expression, as the ratio of array- to sequencing-based experiments was 6 to 1 as of the 2015 publication and the rate of microarray submissions was still growing (Kolesnikov et al., 2015). Future integrative analyses of gene expression will require facing the challenge of combining these data types; a task of utmost importance for rare diseases or understudied biological processes and organisms where all available assays will be required to discover robust signatures or biomarkers. Thus, effective strategies for combining data from the two platforms—perhaps in a manner that leverages both the advantages of RNA-seq and the abundance of microarray data—are paramount to transcriptomic and functional genomic experiments going forward.

Much work has been performed to develop methods for effectively combining multiple cohorts or batches of gene expression data (Johnson et al., 2007;Leek and Storey, 2007;Gagnon-Bartsch and Speed, 2012;Sweeney et al., 2016), but no method has been widely adopted for the problem of combining mixed platform data. Quantile normalization (QN) is a widely used normalization technique originally utilized for microarray data (Bolstad et al., 2003), but it has also been adopted for RNA-seq data normalization (Law et al., 2014) and in some cases, cross-platform normalization (Li et al., 2015). Probe Region Expression estimation Based on Sequencing (PREBS) was developed to make RNA-seq and microarray data more comparable (Uziela and Honkela, 2015), but this method requires raw reads and probe specific information and may not be feasible for large-scale public data efforts. Training Distribution Matching (TDM) was developed by our group to make RNA-seq data more comparable to microarray data from transcript abundances specifically for machine learning applications (Thompson et al., 2016). In that work, it was demonstrated that QN, TDM and a method from the analysis of graphs, nonparanormal normalization (NPN), had good performance in the supervised learning evaluation (Thompson et al., 2016). However, combining platforms was not evaluated.

Here, we present a series of experiments to test what normalization approaches can be used to combine microarray and RNA-seq data for multiple applications: supervised machine learning, unsupervised machine learning, and differential expression analysis. We specifically add varying amounts of RNA-seq data to our training sets to assess at what point performance begins to suffer. We find that QN, TDM, NPN, and standardized scores are all suitable for some use cases, with the widely adopted QN performing well for machine learning applications in particular.

## 2. Methods and Data

We aimed to assess the extent to which it was possible to effectively normalize and combine microarray and RNA-seq data for use as a training set for machine learning applications. We assessed performance on test sets comprised entirely of microarray data and entirely of RNA-seq data. To design such an experiment, we required a data set that had matched samples— sets of samples run on microarray and RNA-seq—and that was of sufficient size.

### 2.1 Evaluation gene expression data

The Cancer Genome Atlas (TGCA) (Cancer Genome Atlas Network, 2012) breast cancer (BRCA) data set includes samples that have been measured with both microarray and RNA-seq platforms. In addition, BRCA has well-defined molecular subtypes that are suitable for use as labels/classes for supervised machine learning approaches we describe below. We used log_2_-transformed, quantile normalized microarray data and RSEM (RNA-seq by Expection Maximization) gene-level count RNA-seq data. (Li and Dewey, 2011) We consider these data to be the products of standard processing pipelines for their respective mRNA expression platforms. For the purpose of these analyses, we restricted the data set to the 520 tumor samples that had been measured on both platforms (termed ‘matched samples’).

### 2.2 Subtype prediction and unsupervised feature extraction

#### 2.2.1 Experimental design

An overview of our experimental design for machine learning evaluations is illustrated in **Fig 1**. Matched samples were split into training (2/3) and test (1/3) sets using the `createDataPartition` function in the caret package (Kuhn, 2012), which takes the balance the class distributions in the training and holdout sets into account (**Fig 1A**). See **Fig S1** for a representative plot of subtype distribution. Two holdout sets were used: a set comprised entirely of RNA-seq data and a set comprised entirely of microarray data. We refer to these as the RNA-seq holdout set and microarray holdout set, respectively. Samples analyzed with RNA-seq were “titrated” into the training set via random selection in 10% increments to produce training sets containing 0%, 10%, 20% … 100% RNA-seq data (**Fig 1B**). The pipeline for partitioning data into training and testing, titration, and normalization was repeated 10 times using different random seeds, as were the downstream analyses (e.g., subtype classification, unsupervised feature construction).

**Figure 1.**
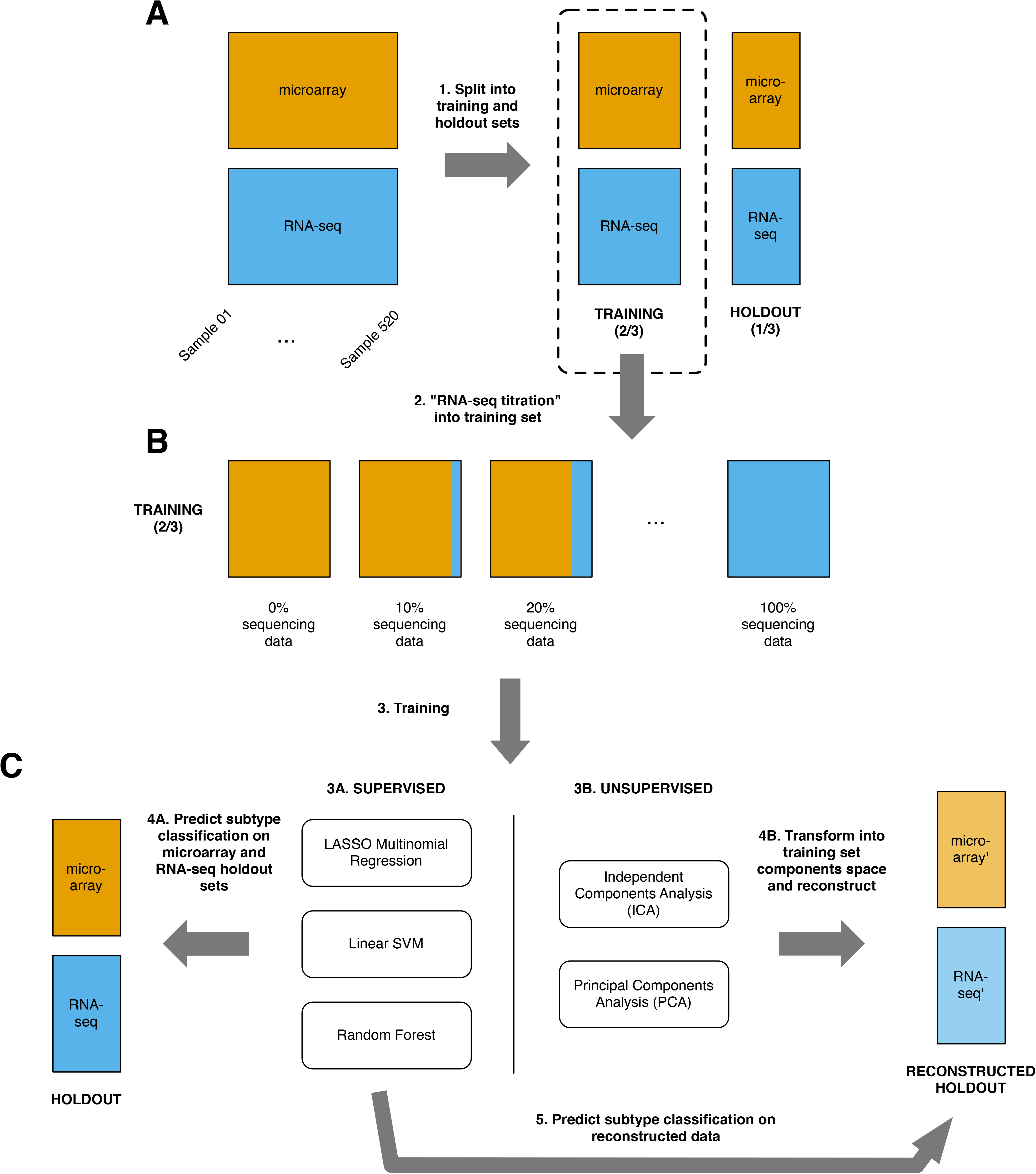
Overview of supervised and unsupervised machine learning experiments. (A) 520 TCGA Breast Cancer samples run on both microarray and RNA-seq were split into a training (2/3) and holdout set (1/3). (B) RNA-seq’d samples were ‘titrated’ into the training set, 10% at a time (0–100%) resulting in eleven training sets for each normalization method. (C) *Machine learning applications.* We used three supervised algorithms to train multi-class (BRCA PAM50 subtype) classifiers on each training set and tested on the microarray and RNA-seq holdout sets. The holdout sets were projected onto and back out of the training set space using two unsupervised techniques, Independent and Principal Components Analysis, to obtain reconstructed holdout sets. The classifiers used in 4A were used to predict on the reconstructed holdout sets.

#### 2.2.2 Cross-platform normalization approaches

For every normalization method, samples source (RNA-seq or microarray) were matched in the training set. In accordance with how these methods would be used in practice, normalization was performed separately for training and holdout sets. For the normalization methods that require a reference distribution (e.g., quantile normalization and Training Distribution Matching), the RNA-seq training and holdout sets were normalized using the microarray data training data as a reference, separately. Only genes measured on both platforms were included; genes that had missing values in the RNA-seq data in all samples (all samples in holdout data, training ‘titration’ samples at any sequencing levels) were removed.

##### Log_2_-transformation (LOG)

As the log_2_-transformed array data contained negative values, the microarray data was inverse log-transformed and then log-transformed such that values are non-negative by adding 1 to each expression value before transformation. All missing values were set to zero. This array data was used in all downstream processing steps.

##### Quantile normalization

QN was performed using the `preProcessCore` R package (Bolstad, 2013). Given the log_2_-transformed microarray distribution (“target”), the `normalize.quantiles.use.target` method will normalize the columns of the RNA-seq data such that the data sets are drawn from the same distribution. For sets entirely comprised of samples on a single platform without an applicable reference, `normalize.quantiles` was used.

##### Training Distribution Matching

TDM was developed specifically to make RNA-seq test data compatible with models trained on microarray data (Thompson et al., 2016); it identifies the relationship that the microarray training data has between the spread of the middle half of the data and the extremes and then transforms the RNA-seq test set such that it has the same relationship between the spread of the middle half of the data and the extremes. We used the TDM R package to perform TDM normalization. For the training set comprised of 0% RNA-seq data, models trained on log_2_-transformed microarray data were used for prediction/reconstruction on the TDM normalized RNA-seq holdout set. TDM was not used when the training set was comprised of 100% RNA-seq data, as there is no relevant microarray data to use as the reference distribution in this case.

##### Non-paranormal normalization

NPN is in essence a rank-transformation followed by placing data where it would fall on a normal distribution (Liu et al., 2009). NPN was performed using the huge R package prior to concatenating samples from both platforms and separately on single-platform holdout sets.

##### Standardizing scores

z-scoring was performed on a per gene basis using the scale function in R prior to concatenating training samples from both platforms. Single-platform holdout sets were z-scored separately. Z-scores or standard scores are calculated 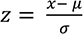, where *μ* and *σ* are the gene mean and standard deviation, respectively

Gene expression values were standardized to the range [0,1] on a per gene basis, either before or after concatenating samples from each platform. Quantile normalization followed by standardizing scores (QN-Z)—used only for differential expression analysis—was performed using the target method as described under *Quantile normalization* above.

#### 2.2.3 Subtype prediction on mixed platform data sets

The PAM50 microarray classifications were used as the subtype labels for supervised analyses (**Fig 1C**). We performed 5-fold cross-validation on training sets for model training and hyperparameter optimization using total accuracy for performance evaluation. We used the Kappa statistic to evaluate performance on holdout data, as classes were not balanced. Briefly, the Kappa statistic takes into account the expected accuracy of a random classifier and is generally considered to be less misleading than observed accuracy alone (Landis and Koch, 1977). We trained the following three classifiers: LASSO logistic regression (Tibshirani, 1996), linear support vector machine (SVM), and random forest. We used the glmnet R package (Friedman et al., 2010) implementation of LASSO. The training of SVM and random forest classifiers was performed using the caret package and utilizing the kernlab (Zeileis et al., 2004) and ranger (Wright and Ziegler, 2015) packages, respectively.

#### 2.2.4 Unsupervised feature extraction from mixed platform data sets

We performed Principal Components Analysis (PCA) and Independent Components Analysis (ICA) on each of the training sets (**Fig 1C**). We performed PCA using the prcomp function in R and ICA using the fastICA function in the fastICA R package (Marchini et al.), setting the number of components to 50. For PCA, we projected the holdout data onto the training data PC space and then reconstructed the holdout data using the first 50 principal components. For ICA, we projected the holdout data onto the training data IC space using the product of the prewhitening and estimated unmixing matrices and reconstructed the holdout data from the projection using the estimated mixing matrix.

We assess reconstruction error (comparing holdout input, *y*, to reconstructed values, *ŷ*) by calculating the mean absolute scaled error (MASE) (Hyndman and Athanasopoulos, 2014). MASE is calculated on a per gene basis as follows:

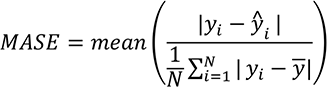

We performed supervised analysis following reconstruction to assess whether the subtype signals were retained or if features were dominated by noise introduced by combining platforms. We used the models trained for subtype classification to predict on the reconstructed holdout sets to assess how well the molecular subtype signal was retained in the reconstructed holdout data (**Fig 1C**). We again used the Kappa statistic to evaluate performance.

### 2.3 Differential expression analyses

We used a standard two-group single-channel experimental design in limma to identify differential expressed genes (Ritchie et al., 2015). We used Benjamini-Hochberg correction (Benjamini and Hochberg, 1995)multiple hypotheses testing, the output of which is known as a false discovery rate (FDR). We used mean-variance modeling at the observational level (VOOM) (Law et al., 2014) as implemented in limma (Ritchie et al., 2015) to pre-process the single platform RNA-seq data. We termed the set of differentially expressed genes (DEGs) as detected at a specified FDR from single platform data microarray and RNA-seq “silver standards” (Sweeney et al., 2017). We again titrated in RNA-seq samples at 10% increments and identified DEGs (**fig 2**). We compared them by calculating the Jaccard similarity, *J*, between the silver standard DEGs, *S*, and the experimental DEGs, *E*, as follows:

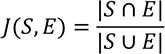

#### 2.3.1 Small sample size experiment

For each number of samples (*n*), we randomly selected *n* (*n* = 3,4,5,6,8,15,25,50) samples each from the Her2 and LumA subtype. To generate platform-specific silver standards, we compared the 2*n* samples in single platform data (100% log_2_-transformed microarray data and 100% RSEM RNA-seq data). For the normalization experiments, we identified DEGs in data sets that were 50% microarray and 50% RNA-seq data. Ten repeats were performed. All genes with an *FDR <* 10% were considered differentially expressed. These sets of DEGs were used to calculate Jaccard similarity.

## 3. Results and Discussion

We performed a series of supervised and unsupervised machine learning evaluations, as well as differential expression analyses, to assess which normalization methods are best suited for combining data from microarray and RNA-seq platforms. We evaluated five normalization approaches for all methods: LOG, NPN, QN, TDM, and standardizing scores (z-scoring; Z).

### 3.1 Non-paranormal normalization, quantile normalization, and Training Distribution Matching allow for training subtype classifiers on mixed platform sets

We trained models to predict BRCA PAM50 subtype on training sets with varying amounts of samples from the RNA-seq platform. We used these models to predict on holdout data sets comprised entirely of microarray data or RNA-seq data (**Fig 1**). We trained three commonly used classifiers: LASSO logistic regression, linear SVM, and random forest. Kappa statistics were used to assess performance. We visualize the Kappa statistics for varying amounts of samples from RNA-seq in the training data in **Fig. 3**. (Note that the pipeline in **Fig 1A-C** was repeated ten times.) The three classifiers showed the same trends across normalization approaches overall, suggesting that the normalization approaches recommended herein will generalize to multiple classification methods.

**Figure 2.**
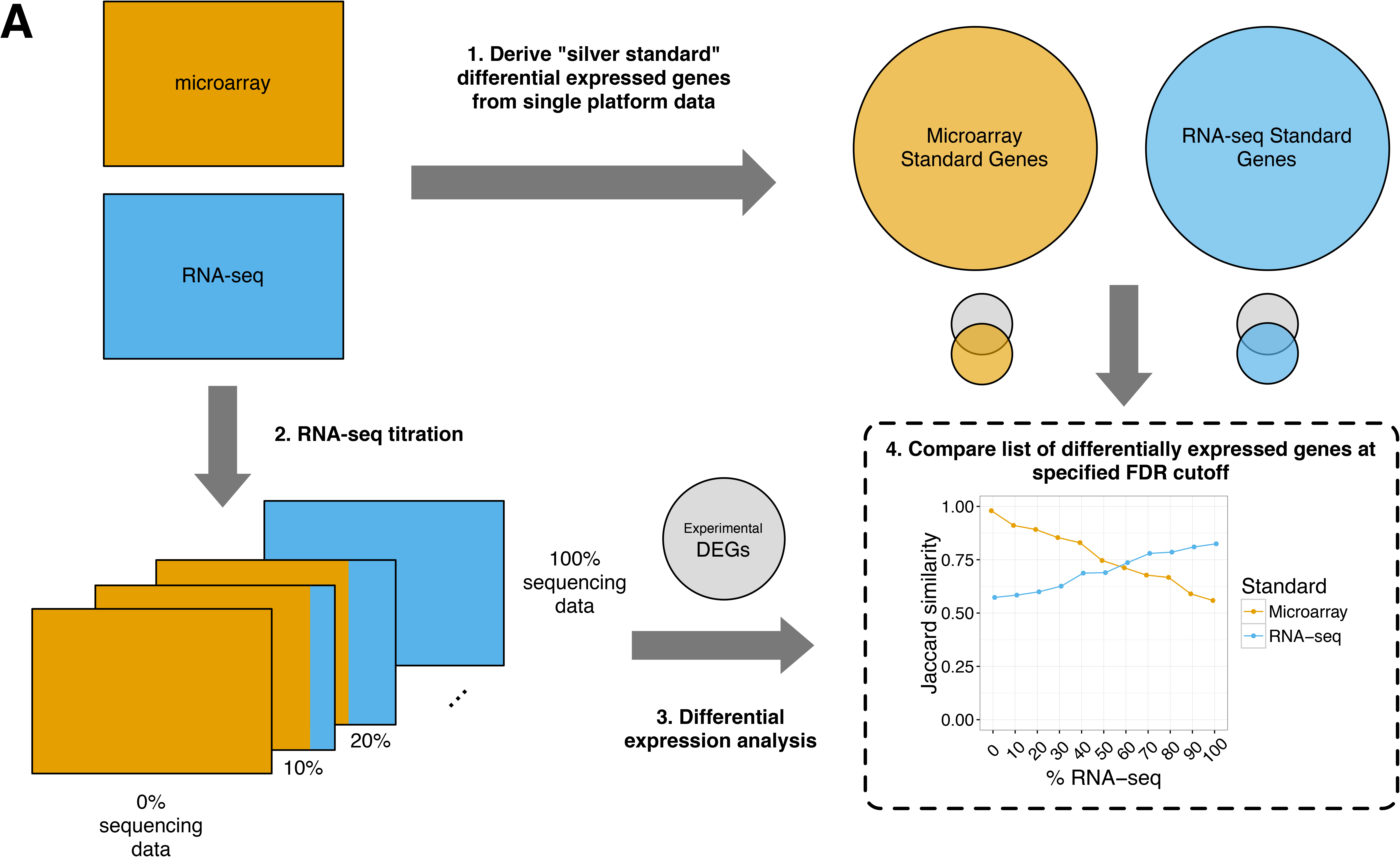
BRCA subtype classifier performance on microarray and RNA-seq holdout data. Violin plots of Kappa statistics from 10 repeats of steps 1-4A from **Figure 1** and for five normalization methods are displayed. Median values are shown as points. (LOG - log2-transformed; NPN – nonparanormal normalization; QN – quantile normalization; TDM – Training Distribution Matching; Z – z-score)

**Figure 3.**
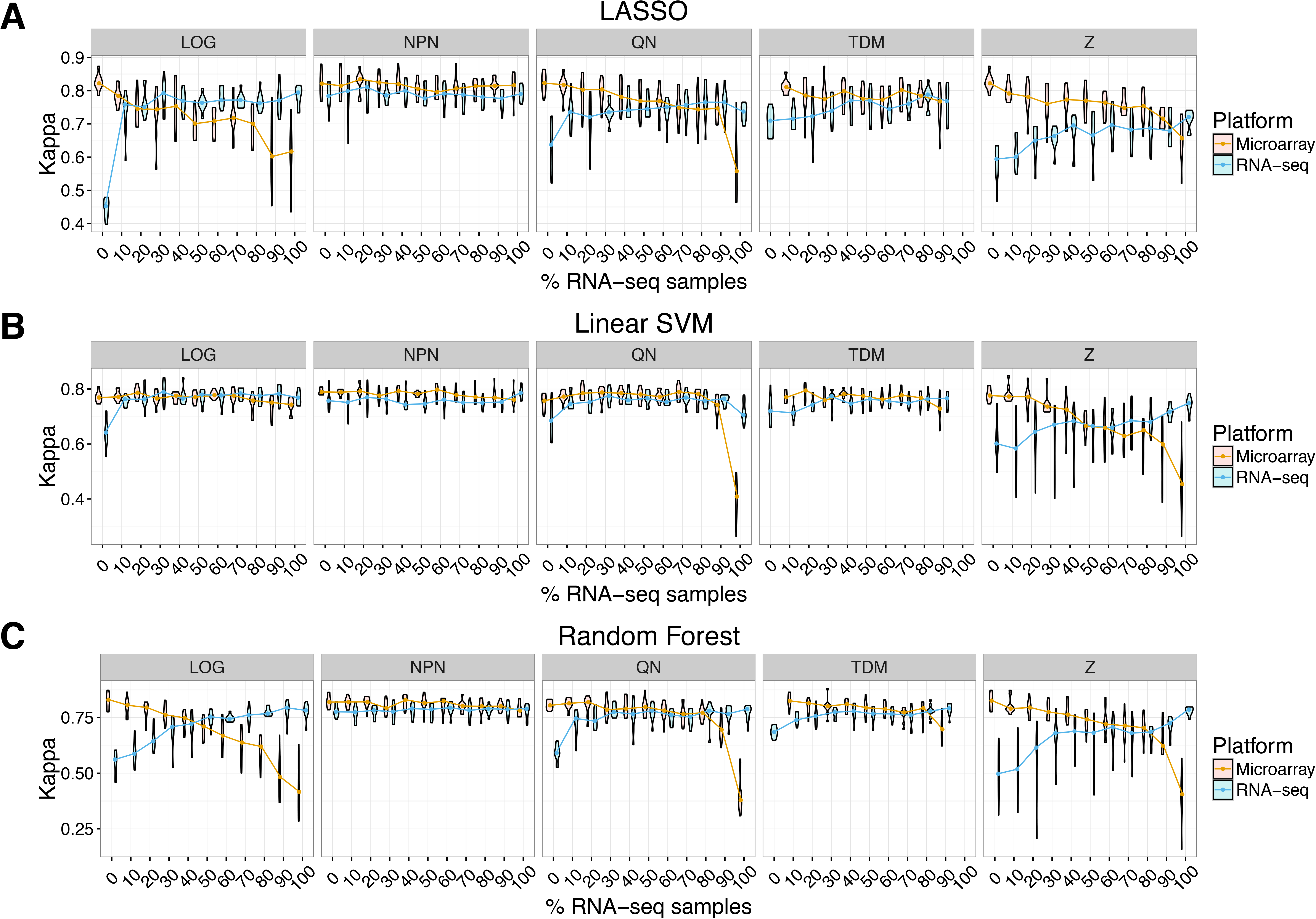
BRCA subtype classifier performance on reconstructed microarray and RNA-seq holdout data. The holdout sets were projected onto the training set Independent or Principal Components reduced dimensional space and back out to obtain reconstructed holdout sets. Subtype prediction on the reconstructed data using the classifiers trained during the supervised analyses was performed. Ten repeats were performed. Violin plots of Kappa statistics are displayed; median values are shown as points. (NPN – nonparanormal normalization; QN – quantile normalization; TDM – Training Distribution Matching; Z – z–score)

Importantly, there were appreciable differences between normalization methods. Log-transformation demonstrated the worst performance. This is expected as we think of this method as a negative control and it was previously shown to be insufficient to make RNA-seq data comparable to microarray (Thompson et al., 2016). We also saw that z-scoring data resulted in the most variable performance. This is not unexpected because the calculation of the standard deviation and mean will be highly dependent on which samples are selected from each platform and the random selection of RNA-seq samples to be included in the training set does not take into account subtype distribution, which may not be known in practice. We found that NPN, QN, and TDM all performed well when moderate (i.e., not extreme) amounts of RNA-seq data are incorporated into the training set. These results are consistent with the high performance of these three methods in our case study of training entirely on microarray data and using solely RNA-seq as a test set (Thompson et al., 2016). These three methods performed well on both the microarray and RNA-seq holdout sets.

Quantile normalization did not perform well at the extremes (0% and 100% RNA-seq data). We attribute this loss of performance to the lack of reference distributions in these cases—for all other amounts of RNA-seq (10 - 90%), the set of microarray data is used as a reference distribution for both the RNA-seq data included in the training set as well as for the holdout set (see Methods). This result reiterates the importance of drawing training and holdout sets from the same distribution, as is well-documented in the machine learning literature, and highlights the necessity of proper cross-platform normalization.

### 3.2 Quantile normalization and Training Distribution Matching are suitable for unsupervised feature extraction from mixed platform data sets

Dimensionality reduction and/or unsupervised feature extraction methods are commonly employed in the analysis of gene expression data. We used two such approaches—PCA and ICA—and evaluated normalization method performance. The molecular subtypes in BRCA are strong, linear signals that we should be able to predict in the holdout sets given the performance of the classifiers visualized in **Fig 3** and should be readily extractable using ICA and PCA. We aimed to identify which normalization methods were most suitable for feature extraction in data sets comprised of a mixture of microarray and RNA-seq data. Our approach was as follows: PCA or ICA was performed on the training sets and then the holdout sets were projected onto the training space and then reconstructed to obtain “reconstructed holdout sets” (see Methods, **Fig 1**). We evaluated performance in two ways: 1) we performed BRCA subtype prediction on the reconstructed sets using the classifiers trained in the supervised analyses and 2) we calculated reconstruction error post-transformation (MASE; see Methods). The Kappa statistics from the first evaluation are visualized in **Fig 4** (see **Fig S2** for performance on LOG normalization data).

**Figure 4.**
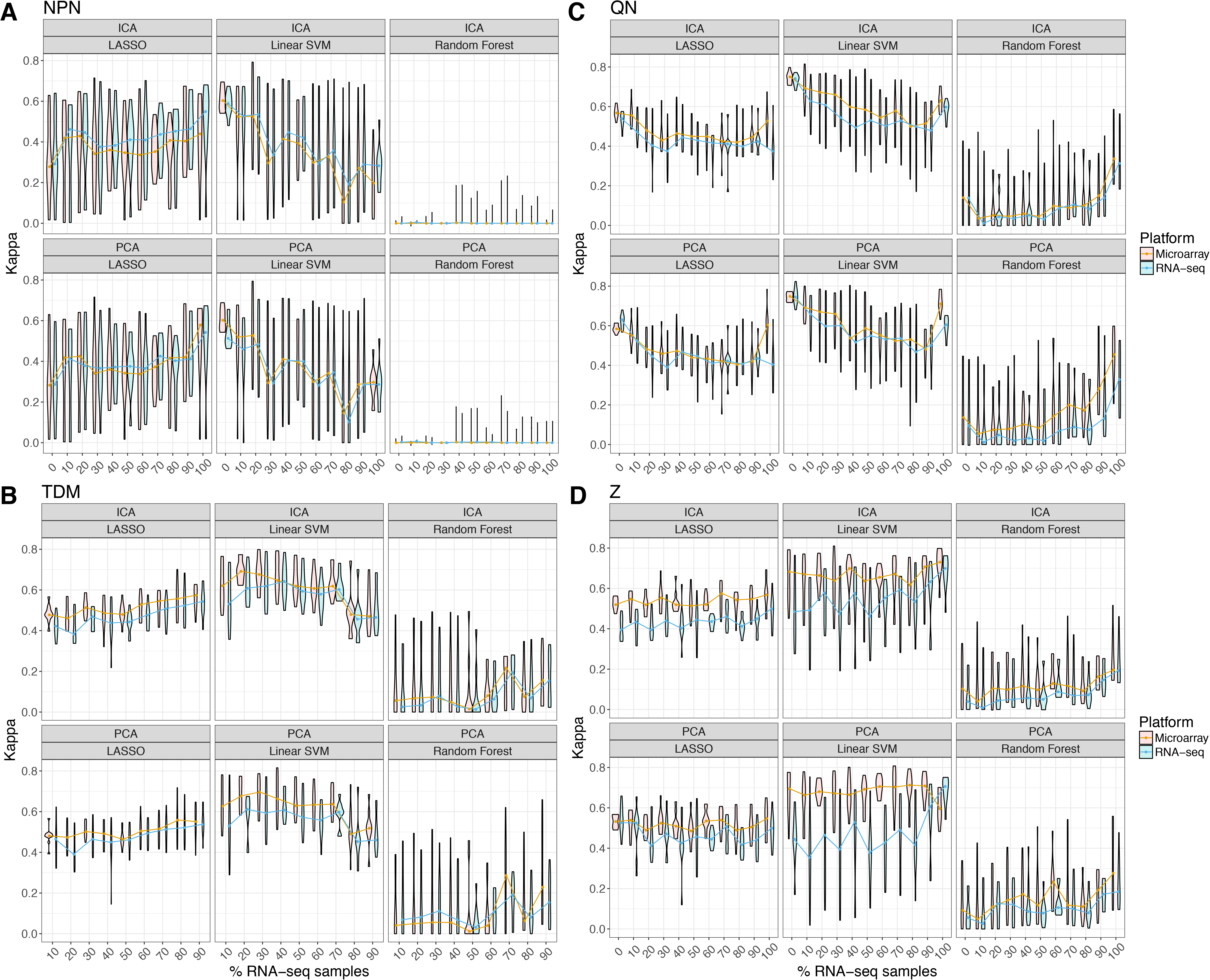
Overview of differential expression experiment. All matched TCGA breast cancer samples (n = 520) were considered when building the platform-specific “silver standards.” These standards are the genes that were differentially expressed at a specified False Discovery Rate (FDR) using data sets comprised entirely of one platform and processed in a standard way: log_2_-transformed microarray data and “untransformed” RSEM count data (preprocessed using the *voom* function in limma). RNA-seq’d samples were ‘titrated’ into the data set, 10% at a time (0–100%) resulting in eleven experimental sets for each normalization method. Differentially expressed genes (DEGs) were identified using the limma package. Lists of experimental DEGs were compared to standard gene sets using Jaccard similarity.

Although we found similar trends among the ICA and PCA reconstructions, we observed differences in classifiers and normalization methods as measured by the Kappa statistics (**Fig 4**). This suggests that ICA and PCA are largely comparable in this particular evaluation. The SVM performance was most robust to reconstruction (**Fig 4**), consistent with expectations for linearly separable class problems. In general, the random forest classifier suffered the largest loss of performance, likely due to gene expression thresholds (rules) used for prediction. In the case of NPN, the near zero random forest Kappa statistics (**Fig 4A**) resulted from predictions of only one class label. We observed the largest differences in performance between the two platform holdout sets with Z normalization (**Fig 4D**). We found that projecting the holdout sets onto QN and TDM normalized training space results in less loss of subtype classifier performance (**Fig 4B-C**). In addition, we observed QN resulted in low reconstruction error (**Fig S3**). This suggests that these two methods are suitable for normalizing sets comprised of data from both platforms for use with unsupervised feature extraction applications.

### 3.3 Differential expression analysis is possible on mixed platform data sets

The identification of genes that are down- or up-regulated between conditions is an exceptionally common analysis performed on gene expression data. Differential expression analysis is also a distinct problem from the supervised and unsupervised machine learning approaches investigated above. For example, prediction of BRCA subtype does not require all genes measured (as demonstrated by the excellent performance of LASSO) and random gene sets are prognostic in BRCA (Venet et al., 2011), but the goal of differential expression analysis in this context is to identify which genes differ between conditions from genome-scale measurements. We sought to identify normalization methods that yielded similar results between data sets comprised of solely one platform and those comprised of a combination of microarray and RNA-seq data (see Methods, **Fig 2**). We termed the set of DEGs as detected at a specified FDR from single platform data microarray and RNA-seq “silver standards” (Sweeney et al., 2017) For these evaluations, we included two additional normalization approaches: untransformed (UN), where untransformed count data (RSEM in this case) and log-transformed microarray data were concatenated together and quantile normalized followed by standardizing scores (QN-Z).

#### 3.3.1 All data

We identified differentially expressed genes between the Her2 and LumA subtypes using the `limma` R package in 100% microarray data and 100% RNA-seq data and identified genes with an FDR *< 5%* to obtain the platform-specific silver standards (see **Fig S4** for proportion of genes identified as differentially expressed at this FDR). We then performed the same comparison in data sets with increasing amounts of RNA-seq samples included. We evaluate the set of DEGs (*FDR <* 5%) from this experiment to both platform-specific standards by computing the Jaccard similarity between the experimental set and the standards. The results of this analysis are visualized in **Fig 5A**. Notably, the microarray and RNA-seq standards have a Jaccard similarity of 0.61 (**Fig 5A**, **LOG panel**), which indicates that the majority of DEGs detected using standard pipelines for each of the platforms are shared but differences exist.

**Figure 5.**
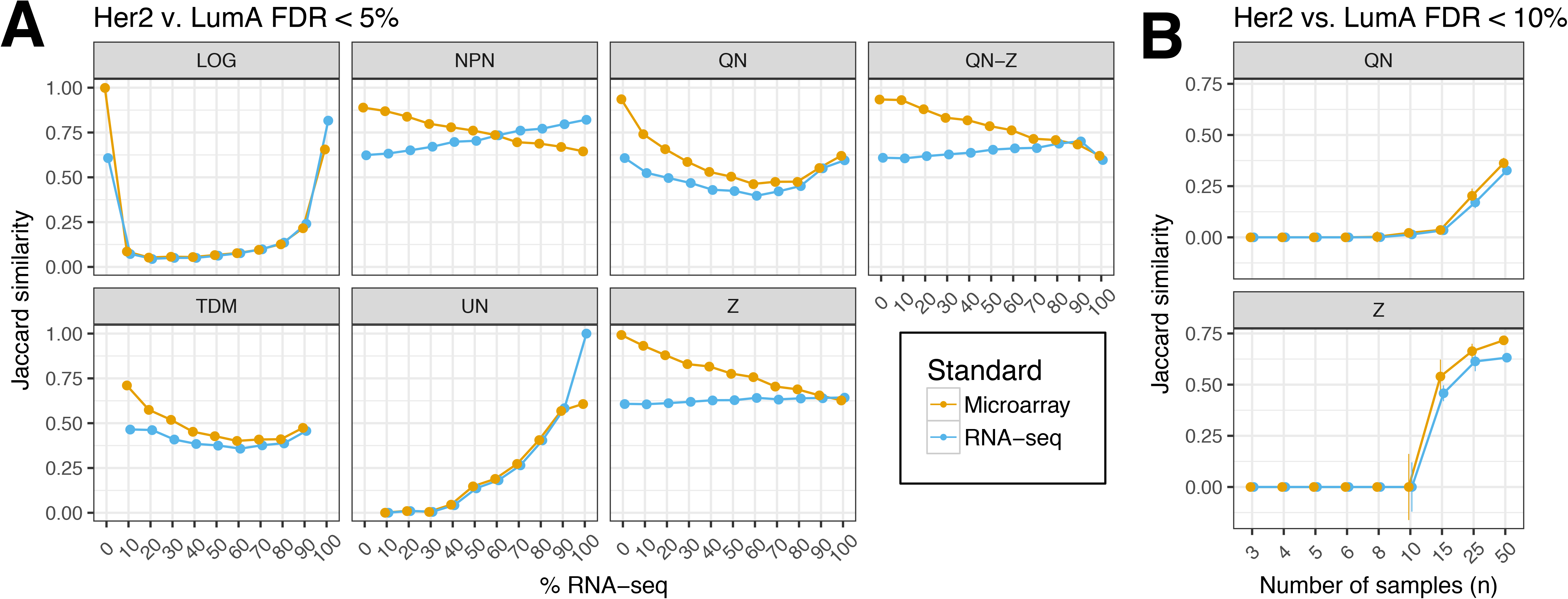
Overlap between platform-specific silver standard differentially expressed genes (DEGs) and experimental DEGs when testing (A) all samples and (B) smaller sample sizes. Genes differentially expressed between Her2 and LumA samples were identified. (A) All Her2 and LumA samples were compared for all normalization methods with varying amounts of RNA-seq data included. All genes with an FDR < 5% were considered differentially expressed. These lists were used to calculate Jaccard similarity. (B) *Small n experiment.* For each number of samples (n), we randomly selected *n* samples each from the Her2 and LumA subtype. To generate platform-specific standards, we compared the 2*n* samples in single platform data (100% log_2_-transformed microarray data and 100% RSEM RNA-seq data). For the normalization experiments, we identified DEGs in data sets that were 50% microarray and 50% RNA-seq data. Ten repeats were performed. All genes with an FDR < 10% were considered differentially expressed. These lists were used to calculate Jaccard similarity. The median and 95% confidence on the median” (+/−1.58 IQR/sqrt(n) Chambers, et al. 1983) of the Jaccard similarity from ten repeats is displayed. (LOG - log2-transformed; NPN – nonparanormal normalization; QN – quantile normalization; QN-Z - quantile normalization followed by z-score; TDM – Training Distribution Matching; UN – untransformed (count data); Z – z-score)

There are marked differences in normalization approach performance as measured by Jaccard similarity to silver standards, with the results mirroring our machine learning assessments. We found that adding log-transformed and untransformed (count) RNA-seq data to the data set, which we regard as negative controls, recovered the fewest silver standard DEGs (for either platform) as demonstrated by the low Jaccard similarity to standards (**Fig 5A**). We found that TDM and QN have similar performance, recovering less than half of the standard DEGs for moderate amounts of RNA-seq data in the data set used to identify DEGs. Using NPN, Z, or QN-Z resulted in detection of the majority of silver standard DEGs, suggesting that these methods are best suited to the differential expression problem. We also performed a similar analysis comparing Basal samples to all other subtypes and observed similar performance across methods (**Fig S5**).

#### 3.3.2 Small sample sizes

The differential expression analyses we describe above utilized all matched BRCA samples. To provide some guidance in combining the two platforms in considerably smaller data sets, we devised a “small n” experiment. We elected to evaluate the QN and Z normalization approaches because of their simplicity and wide usage. We used varying sample sizes from 3 to 50 samples (see Methods, **Fig 5B**). We again used Jaccard similarity to single platform standards to evaluate normalization method performance.

We hypothesized that the z-score approach would only perform well with sufficient *n* because a larger sample size would be required to reliably calculate the gene means and standard deviations. Our results support this hypothesis (**Fig 5B**), as we only observe recovery of DEGs in the mixed platform data sets when *n* ≥ 15. We observed poor performance, regardless of sample size, with QN (**Fig 5B**). This is perhaps unsurprising given the performance in the larger data set (**Fig 5A**). In total, these results suggest that caution is required in when combining multiple platforms *particularly* when sample sizes are small.

## 4. Conclusions

Herein we perform, to our knowledge, the first examination of cross-platform normalization for machine learning on training sets comprised of data measured on both popular gene expression platforms—RNA-seq and microarray—and demonstrate that it is possible to combine these data types for use with supervised and unsupervised applications as well as differential expression analysis. We find that QN and TDM perform well for both types of machine learning approaches. NPN and Z are also appropriate for some applications, but care may be required in of z-scoring the data to ensure that that the distributions are comparable (e.g., distributions of classes, which may not be known). We also find that combining platforms is not appropriate for data sets with a small number of replicates (**Fig 5B**).

This study has some important limitations. Because of our experimental design, we required a large data set of samples run on both platforms. This limited our work to one data set of high quality: TCGA BRCA data. BRCA subtypes are well-defined signatures that have evident linear expression patterns. As a result, our guidance may not generalize to nonlinear classifiers, data sets of poor quality, or small sample sizes.

Nevertheless, this work indicates that it is possible to perform model training and differential expression analysis on microarray and RNA-seq. Combining both platforms could allow models to take advantage of the additional information captured in some RNA-seq experiments while benefiting from the substantially greater abundance of microarray data. Training sets comprised of samples run on both platforms will be a new realty as RNA-seq becomes the platform of choice and the ability to perform such analyses will be of particular importance for understudied biological problems.

## Acknowledgements

We thank Gregory Way for helpful code review and Amy Campbell for proof-reading of the manuscript.

